# Calcium-independent lipid release from astrocytes modulates neuronal excitability

**DOI:** 10.1101/2020.01.12.903393

**Authors:** Fushun Wang, Heather B. Bradshaw, Salvador Pena, Beata Jablonska, Julia Xavier, Siddharth Chittaranjan, Sheng Gong, Baoman Li, Devin Chandler-Militello, Lane K. Bekar, Nathan A. Smith

**Affiliations:** Departments of Neurosurgery, University of Rochester School of Medicine and Dentistry, Rochester, NY 14642, USA; Institute of Brain and Psychological Sciences, Sichuan Normal University, Chengdu, Sichuan 610060, China; Division of Glial Disease and Therapeutics, Center for Translational Neuromedicine, Department of Neurosurgery, University of Rochester Medical Center, Rochester, NY 14642, USA.; Center for Translational Neuromedicine and the Department of Neurology, University of Rochester Medical Center, Rochester, NY 14642, USA; Department of Psychological and Brain Sciences, Indiana University, Bloomington, IN 47405, USA; Department of Anesthesiology, Wake Forest School of Medicine, Winston-Salem, North Carolina; Center for Neuroscience Research, Children’s National Research Institute, Children’s National Hospital, Washington DC 20010, USA; George Washington University, Washington DC 20052, USA; Department of Physiology and Biophysics, Case Western Reserve University School of Medicine, Cleveland, OH 44106, USA; Department of Anatomy, Physiology, and Pharmacology, University of Saskatchewan, Saskatoon, SK S7N 5E5; Del Monte Institute for Neuroscience, Department of Neuroscience, University of Rochester, School of Medicine and Dentistry, Rochester, USA

## Abstract

An accumulating amount of data suggests that Ca^2+^-dependent gliotransmitter release plays a key role in the modulation of neuronal networks. Here, we tested the hypothesis that in response to agonist exposure, astrocytes release lipid modulators through activation of Ca^2+^-independent phospholipase A_2_ (iPLA_2_) activity. We found that cultured rat astrocytes treated with selective ATP and glutamatergic agonists released arachidonic acid (AA) and/or its derivatives, including the endogenous cannabinoid 2-arachidonoyl-sn-glycerol (2AG) and prostaglandin E2 (PGE_2_). Surprisingly, the buffering of cytosolic Ca^2+^ resulted in a sharp increase in agonist-induced lipid release by astrocytes. In addition, the astrocytic release of PGE_2_ increased miniature excitatory postsynaptic potentials (mEPSPs) by inhibiting the opening of neuronal Kv channels in brain slices. This study provides the first evidence showing that a Ca^2+^-independent pathway regulates the release of PGE_2_ from astrocytes and further demonstrates the functional role of astrocytic lipid release in the modulation of synaptic activity.

**SIGNIFICANCE:** Until now, most studies that implicate astrocytes in the modulation of synaptic activity have focused on Ca^2+^-dependent release of traditional gliotransmitters such as D-serine, ATP, and glutamate. Mobilization of intracellular stores of Ca^2+^ occurs within a matter of seconds, but this novel Ca^2+^-independent lipid pathway in astrocytes could occur on a faster time scale and thus play a role in the rapid signaling processes involved in synaptic potentiation, attention, and neurovascular coupling.

## INTRODUCTION

In addition to being a major contributor to the dry weight of the adult brain, lipids are an essential component of the phospholipid bilayer and are mainly composed of long-chain polyunsaturated fatty acids (PUFAs), such as arachidonic acid (AA) and docosahexaenoic acid (DHA) (Sinclair, 1975). The liver is the major site of AA synthesis, but the brain can produce AA and DHA in situ from their precursor fatty acids, linoleic and linolenic acids (Dhopeshwarkar and Subramanian, 1976). Astrocytes play a central role in the synthesis of AA and DHA in the brain. Through their vascular endfeet, astrocytes have prime access to fatty acid precursors that arrive by crossing the blood‒brain barrier (BBB) and serve as the major site for processing essential fatty acids in the central nervous system (CNS) (Moore, 1993). Astrocytes also play a key role in the macroscopic distribution of lipids in the brain parenchyma via perivascular glymphatic flux (Rangroo Thrane et al., 2013a; Plog and Nedergaard, 2018).

Lipids have gained much attention for their role as bioactive mediators in the CNS (Carta et al., 2014; Ledo et al., 2019). Numerous studies have focused on lipids in relation to functional hyperemia and synaptic activity. For instance, PGE_2_ is a potent vasodilator and vasoconstrictor that regulates CNS blood flow (Zonta et al., 2003; Takano et al., 2006; Gordon et al., 2007; Dabertrand et al., 2013; MacVicar and Newman, 2015; Czigler et al., 2019) and modulates the membrane excitability of CA1 pyramidal neurons during synaptic activity (Chen and Bazan, 2005). Furthermore, AA and its derivatives are important intracellular secondary messengers that can modulate the activities of various ion channels (Piomelli, 1993; Meves, 1994; Horimoto et al., 1997; Boland and Drzewiecki, 2008; Cordero-Morales and Vasquez, 2018). In addition, PGE_2_ can suppress the outwardly rectifying Kv current in sensory neurons (Nicol et al., 1997; Evans et al., 1999), whereas AA suppresses Kv channels in the soma or dendrites of pyramidal neurons and consequently broadens their presynaptic action potentials (Carta et al., 2014) and enhances EPSPs (Ramakers and Storm, 2002). However, those studies focused on the effects of neuronal lipid release on synaptic activity and paid little attention to receptor-mediated pathways by which astrocytic lipids might influence synaptic activity. Given that the majority of AA, DHA, and other lipids present in the extracellular fluid are produced by astrocytes (Moore et al., 1991), receptor-mediated release could be a significant factor in the modulation of synaptic activity.

In culture, astrocytes can release AA via a cPLA_2_ Ca^2+^-dependent pathway upon the activation of metabotropic glutamate (mGluR) and P2Y purine (ATP) receptors (Bruner and Murphy, 1990; Stella et al., 1994; Stella et al., 1997; Chen and Chen, 1998). In addition, astrocytes also express the Ca^2+^-independent PLA_2_ (iPLA_2_) enzyme (Sun et al., 2005). Both of these isoforms are activated by the G-protein βγ subunit (Jelsema and Axelrod, 1987; Murayama et al., 1990; van Tol-Steye et al., 1999), but iPLA2 does not require Ca^2+^ or PKC phosphorylation for activation (Winstead et al., 2000). Receptor-stimulated iPLA_2_ activation can induce the release of AA and DHA by numerous types of cells (Gross et al., 1993; Portilla et al., 1994; Akiba et al., 1998; Seegers et al., 2002; Tay and Melendez, 2004), but this activation has not been fully explored in the case of astrocytes.

Previous work in our laboratory and elsewhere has demonstrated that astrocytes are capable of releasing gliotransmitters upon stimulation with mGluR or ATP receptor agonists, but the impact of this signaling pathway on synaptic activity remains controversial. Recent studies with genetically encoded calcium indicators allowed for the identification of localized Ca^2+^ signals within astrocytic fine processes and confirmed Ca^2+^-dependent astrocytic effects on synaptic activity (Yu et al., 2018). However, the existence of Ca^2+^-independent lipid signaling in astrocytes is still unclear. Therefore, in this study, we tested the hypothesis that astrocytes support Ca^2+^-independent lipid signaling to modulate synaptic transmission. We used Ca^2+^ chelation to show that ATP and mGluR agonists can release lipids through iPLA_2_ activation, resulting in the potentiation of synaptic activity. Because we currently lack tools to assess Ca^2+^-independent signaling in astrocyte fine processes adjacent to synapses, we used Ca^2+^ inhibition on a large scale to evaluate this phenomenon (see discussion).

## MATERIALS AND METHODS

### Culture

Cultured neocortical astrocytes were prepared from postnatal day 1 or 2 Wistar rat pups (Taconic Farms, Inc.) of either sex, as previously described (Lin et al., 1998). In brief, cerebral cortices were dissected on ice, and the meninges were removed. The tissue was washed three times in Ca^2+^-free Hanks’ balanced salt solution, triturated, filtered through a 70 µm nylon mesh, and centrifuged. The pellet was resuspended in 10% fetal bovine serum in Dulbecco’s modified Eagle’s medium (DMEM)/F12 containing penicillin (100 IU ml^-1^) and streptomycin (100 µg ml^-1^) and transferred to culture flasks. The cells were maintained at 37 °C in an incubator with humidified air and 5% CO_2_. The medium was changed after 24 hours and twice a week thereafter. More than 95% of the cells immunostained were GFAP positive. When the cells became confluent, they were rinsed two times in Ca^2+^-free Hanks’ balanced salt solution, suspended in 0.05% trypsin-containing PBS for 1 minute, resuspended in DMEM/F12, centrifuged to remove the trypsin-containing supernatant, and then plated in 24-well plates. The experiments were performed when the cells were 95% confluent.

### Viral vectors and viral transduction

Viral vectors driving GFAP cyto-GCaMP6f (Baljit S. Khakh) were obtained from the University of Pennsylvania Vector Core (AAV 2/5 serotype). Secondary rat astrocytic cultures were transduced with the AAV GFAP cyto-GCaMP6f. After transduction, the cultures were incubated at 37 °C for 5 days prior to Ca^2+^ imaging experiments.

### Ca^2+^ imaging

Cultured cells in 24-well plates were transduced with the AAV GFAP cyto-GCaMP6f and incubated with various pharmacological agents for 30 minutes at 37 °C. Using confocal microscopy (Olympus FV500), calcium wave activity was evaluated by adding an equal volume of medium containing 100 µM ATP to each well. Relative increases in the fluorescence signal evoked by P2Y receptor agonist exposure were compared with baseline fluorescence (ΔF/F) as previously described (Nedergaard, 1994; Smith et al., 2018).

### Radiolabeling and assessment of AA release

Confluent rat astrocytic cultures were incubated with 100 nCi [5,6,8,9,11,12,14,15-^3^H]-AA (PerkinElmer) overnight before experiments. The cells were washed three times with serum-free medium and then allowed to recover for 20 minutes. Before stimulating P2YRs with 100 µM ATP, the cells were incubated with the appropriate inhibitors for 10 to 12 minutes. Aliquots of medium were taken 15 minutes after agonist stimulation, and the levels of ^3^H-AA and/or metabolites were measured by liquid scintillation counting.

### HPLC/MS/MS analysis of lipids

Confluent rat astrocytes were washed with serum-free media and then allowed to recover for 1 hour in serum-free media. At this time, the cells were incubated with vehicle or 20 µM cyclopiazonic acid (CPA) for 15 minutes and then challenged with 100 µM ATP or vehicle for 15 minutes, as described in the section on [^3^H]-AA experiments. The media were then removed and put aside, and 2 ml of HPLC-grade methanol was added to the flask for 5 minutes. After removing the methanol, an additional 2 ml of HPLC-grade methanol was added, the cells were scraped from the sides of the flask, and the contents were added to the previous two retained fractions. Deuterium standards were added to a final concentration of 200 pM, and the samples were centrifuged at 19,000 × g and 24 °C. Lipids in the eluent were partially purified on 500 mg C18 Bond Elut solid phase extraction columns (Varian), concentrated into fractions of 40, 60, 85, and 100% methanol, and analyzed using HPLC/MS/MS (Shimadzu SCL10Avp-Wilmington, DE, USA; API 3000 Applied Biosystems/MDS SCIEX, Foster City, CA, USA) as previously described (Leishman et al., 2018). Over 50 lipids were targeted in these analyses, including 40 lipoamines, 3 acylglycerols, 2 free fatty acids, and 2 prostaglandins (namely, X and Y). The results of pilot studies showed that the analyte concentrations were too low for reliable detection in the medium alone, but the extraction of medium and cells together provided sufficient lipid concentrations for reliable assays.

### PGE_2_ release assessment via PGE_2_ immunoassay

Confluent rat astrocytic cultures were washed 3 times with serum-free medium and then allowed to recover for 20 minutes. Before stimulation with 100 µM ATP, the cells were incubated with the appropriate P2Y inhibitors for 10 to 12 minutes. Aliquots of the medium were collected 15 minutes after stimulation and PGE2 content was determined using an immunoassay kit (Cayman Chemicals) according to the manufacturer’s instructions.

### Western blotting

Protein in samples harvested from the 24-well plates was separated by SDS‒PAGE and transferred to a nitrocellulose membrane, which was then blocked with Tris-buffered saline containing 0.05% (wt/vol) Tween 20 and 5% nonfat dry milk. The primary antibodies used were anti-iPLA2 (Sigma, St. Louis, MO) and anti-β-actin (Cell Signaling, Danvers, MA) at 1:1000 to 1:2000 dilutions in blocking buffer. Chemiluminescence from horseradish peroxidase-linked secondary antibodies was detected using the ChemiDoc™ XRS+ System and running Image Lab™ software.

### Isolation of human fetal astrocytes

Human fetal forebrain tissues were obtained from second-trimester aborted fetuses at 20 weeks of gestational age. Tissues were obtained from aborted fetuses, with informed consent and tissue donation approval from the mothers using protocols approved by the Research Subjects Review Board of the University of Rochester Medical Center. No patient identifiers were made available to or known by the investigators; no samples with known karyotypic abnormalities were included. Forebrain tissue samples were collected and washed 2-3 times with sterile Hank’s balanced salt solution containing Ca^2+^/Mg^2+^ (HBSS^+/+^). The cortical plate region (CTX) of the fetal forebrain was dissected and separated from the ventricular zone/subventricular zone (VZ/SVZ) portion. CTX was then dissociated from papain as previously described (Keyoung et al., 2001). The cells were resuspended at a density of 2-4 x 10^6^ cells/ml in DMEM/F12 supplemented with N2, 0.5% FBS, and 10 ng/ml bFGF and plated in suspension culture dishes. The day after dissociation, the cortical cells were recovered and subjected to magnetic activated cell sorting (MACS) to purify the astrocyte progenitor population. The recovered cells were briefly incubated with CD44 microbeads according to the manufacturer’s recommendations (Miltenyi Biotech). The cells were then washed, resuspended in Miltenyi Washing buffer, and bound to a magnetic column (Miltenyi Biotech). The bound CD44+ astrocyte progenitor cells were eluted, collected, and then washed with DMEM/F12. The purified human fetal astrocyte progenitors were cultured in DMEM/F12 supplemented with N2 and 5% FBS to further differentiate them. To prepare culture dishes for PGE2 immunoassays or immunocytochemistry, fetal cortical astrocytes were dissociated with TrypLE (Invitrogen) into single cells and then plated onto poly-*L*-ornithine/laminin-coated 24-well plates (50,000 cells per well).

### shRNA-mediated lentiviral knockdown of iPLA_2_ in astrocytes

Rat astrocyte cultures were plated in a 24-well plate and grown to approximately 50% confluence. The cultures were transduced overnight by adding either group VI iPLA_2_ shRNA (r) lentiviral particles (sc-270117-V) or control shRNA lentiviral particles-A (sc-108080) directly to culture medium containing polybrene (sc-134220) according to the manufacturer’s instructions with minor modifications (all from Santa Cruz Biotechnology, Santa Cruz). At 24 hours after transfection, the culture medium was removed, and fresh culture medium without polybrene was added. Experiments and Western blot analysis were performed 7 days after transduction.

### Acute hippocampal slice preparation and electrophysiology

Unless otherwise noted, 15–21-day-old C57BL/6 (Charles River, Wilmington, MA), MrgA1^+/-^ transgenic, and littermate control MrgA1^-/-^ pups (courtesy of Dr. Ken McCarthy) (Fiacco et al., 2007) of either sex were used to prepare hippocampal slices as previously described (Wang et al., 2012). The pups were anesthetized in a closed chamber with isoflurane (1.5%) and decapitated. The brains were rapidly removed and immersed in an ice-cold cutting solution that contained (in mM) 230 sucrose, 2.5 KCl, 0.5 CaCl_2_, 10 MgCl_2_, 26 NaHCO_3_, 1.25 NaH_2_PO_4_, and 10 glucose, at a pH of 7.2-7.4. Coronal slices (400 μm) were cut using a vibratome (Vibratome Company, St. Louis) and transferred to oxygenated artificial cerebrospinal fluid (aCSF) that contained (in mM) 126 NaCl, 4 KCl, 2 CaCl_2_, 1 MgCl_2_, 26 NaHCO_3_, 1.25 NaH_2_PO_4_, and 10 glucose (pH = 7.2-7.4; osmolarity = 310 mOsm). The slices were then incubated in aCSF for 1-5 hours at room temperature before electrophysiological measurement. The experiments were performed at room temperature (21-23 °C). During the recordings, the slices were placed in a perfusion chamber and superfused with aCSF gassed with 5% CO_2_ and 95% O_2_ at room temperature. The cells were visualized with a 40× water-immersion objective and differential inference contrast (DIC) optics (BX51 upright microscope, Olympus Optical, New York, NY). Patch electrodes were fabricated from thin-wall glass filaments (World Precision Instruments) on a vertical puller; the resistance of the pipette was approximately 6-9 MΩ with the addition of intracellular pipette solution. The pipette solution contained 140 mM K-gluconate, 5 mM Na-phosphocreatine, 2 mM MgCl_2_, 10 mM HEPES, 4 mM Mg-ATP, and 0.3 mM Na-GTP (pH adjusted to 7.2 with KOH). The current‒voltage (I‒V) curves of the voltage–gated potassium currents were recorded under a voltage clamp using an AxoPatch MultiClamp 700B amplifier (Axon Instruments, Forster City, CA). When measuring outward currents, QX314 (0.5 mM) was added to the pipette solution to block Na^+^ currents. For recordings of miniature excitatory postsynaptic potentials (mEPSPs), 0.5 µM TTX was added to the aCSF. The junction potential between the patch pipette and the bath solution was zeroed before the giga seal was formed. Patches with seal resistances less than 1 GΩ were rejected. The data were low-pass filtered at 2 kHz and digitized at 10 kHz with a Digidata 1440 interface controlled by pClamp Software (Molecular Devices, Union City, CA).

### Pharmacological agents used in culture and slice experiments

Adenosine 5 -triphosphate (ATP, 100 µM); cyclopiazonic acid (CPA, 20 µM); *trans*-(1S, 3R)-1-Amino-1,3-cyclopentanedicarboxylic acid (*t*-ACPD, 100 µM); (±)-α-amino-3-hydroxy-5-methylisoxazole-4-propionic acid hydrobromide ((±)-AMPA, 100 µM); Prostaglandin E_2_ (PGE_2_, 50 µM, Tocris); Phe-Met-Arg-Phe amide (FMRF, 15 µM); Thr-Phe-Leu-Leu-Arg-NH_2_ (TFLLR-NH_2_, 30 µM, Tocris); N-acetyl-Asp-Glu (NAAG, 100 µM); (1*R*, 4*R*, 5*S*, 6*R*)-4-amino-2-oxabicyclo [3.1.0] hexane-4,6-dicarboxylic acid disodium salt (LY379268, 100 µM, Tocris); calmidazolium chloride (CMZ, 2 µM, Tocris); methyl arachidonyl fluorophosphonate (MAFP, 10 µM); bromoenol lactone (Bel, 10 µM); AA (50 µM); *N*-(2,6-dimethylphenylcarbamoylmethyl) triethylammonium chloride (QX314, 1 mM, Tocris); 4-(4-,9-diethoxy-1,3-dihydro-1-oxo-2*H*-benz[f]isoindol-2-yl)-*N*-(phenylsulfonyl) benzeneacetamide (GW627368X 3 µM, Tocris); 6-Isopropoxy-9-xanthone-2-carboxylic acid (AH6809, 10 µM, Tocris); *N*-(piperidin-1-yl)-5-(4-iodophenyl)-1-(2,4-dichlorophenyl)-4-methyl-1*H*-pyrazole-3-carboxamide (AM251, 5 µM, Tocris); Tetrodotoxin (TTX, 0.5 µM, Tocris); 1,2-bis(2-aminophenoxy)ethane-*N*,*N*,*N*’,*N*’-tetraacetic acid (BAPTA, 50 µM, Tocris); and 1,2-*Bis*(2-aminophenoxy)ethane-*N*,*N*,*N*’,*N*’-tetra-acetic acid tetrakis (acetoxymethylester) (BAPTA-AM, 20 µM) were utilized. All chemicals were obtained from Sigma unless otherwise noted.

### Statistical analysis

Statistical significance was evaluated by one-way ANOVA and *post hoc* tests (Tukey and Dunn) using Prism software and deemed significant when P<0.05 for the [^3^H]-AA and PGE_2_ assay. The normality of the data was evaluated by the Shapiro‒Wilk test with a = 0.05. For electrophysiology experiments, significance was determined by paired or unpaired t tests or Tukey‒Kramer *post hoc* multiple comparison tests. HPLC/MS/MS lipidomic data were analyzed with ANOVA and Fisher’s LSD *post hoc* test using SPSS when P<0.05 or P<0.10. All results are reported as the mean ± s.e.m.

## RESULTS

### Ca^2+^-independent release of [^3^H]-AA and its metabolites from cultured astrocytes

We first assessed the efficiency of preloading with an inhibitor of the endoplasmic reticulum (ER) Ca^2+^ pump, cyclopiazonic acid (CPA) (20 µM), or the cytosolic Ca^2+^ chelator 1,2-bis(o-aminophenoxy) ethane-N,N,N′,N′-tetraacetic acid (acetoxymethyl ester) (BAPTA-AM) (20 µM), which was able to block ATP (100 µM)-induced increases in cytosolic Ca^2+^ in cultured rat astrocytes. Images of cytosolic Ca^2+^ (AAV GFAP cyto-GCaMP6f) showed that the purine agonist ATP (100 µM) induced a prompt increase in Ca^2+^ that was completely blocked in CPA- and BAPTA-loaded cultures, whereas 10 µM methylarachidonyl fluorophosphate (MAFP), a nonspecific inhibitor of both cPLA_2_ and iPLA_2_, or 10 µM bromoenol lactone (Bel) (Cornell-Bell et al.), a specific inhibitor of iPLA_2_, did not affect ATP-induced increases in Ca^2+^ in astrocytic culture **(Figure 1A-C)**.

**Figure 1:**
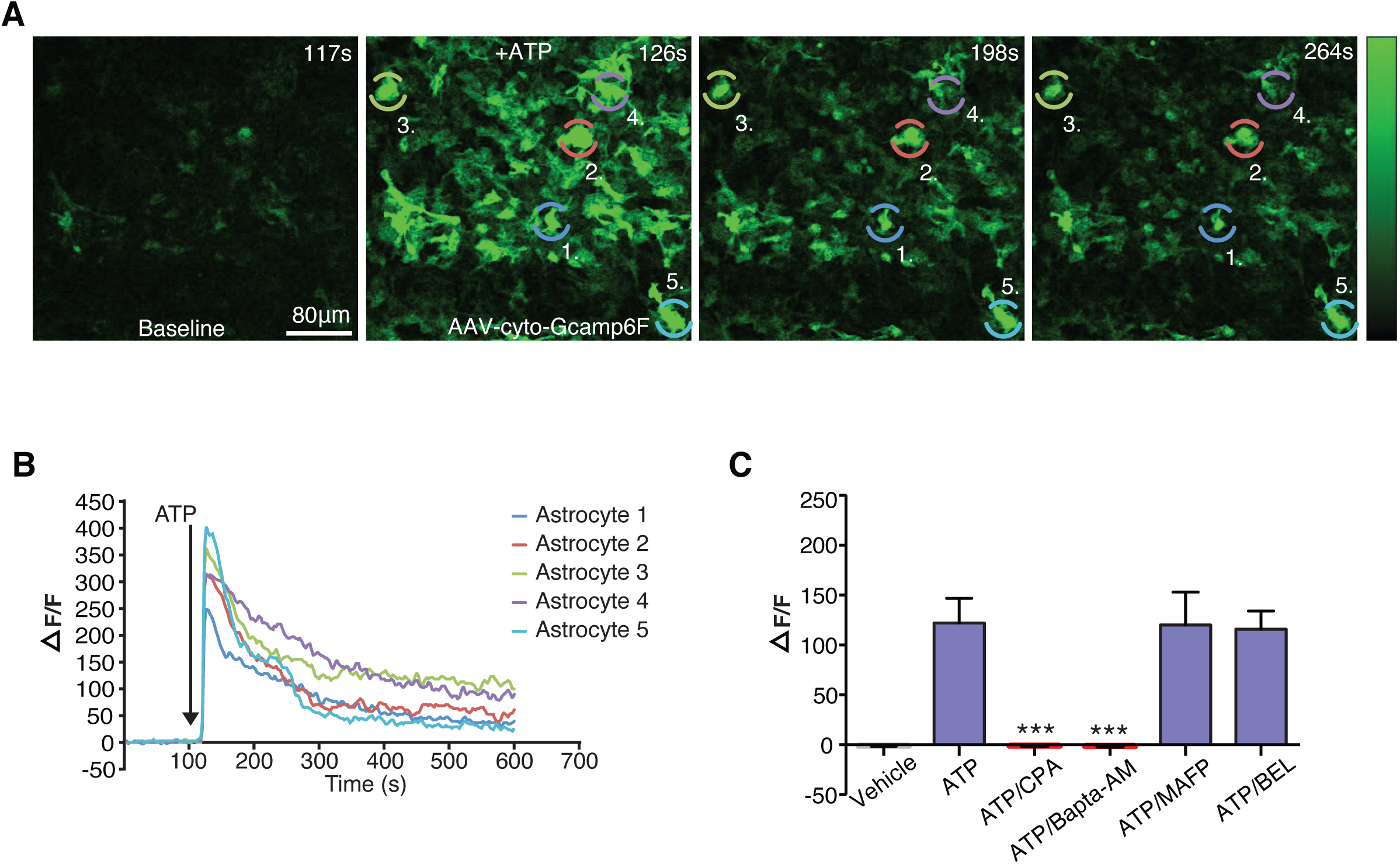
GPCR-evoked Ca^2+^ transients in astrocytic cultures. A.) Cultured astrocytes were transduced with the AAV GFAP cyto-GCaMP6f, and the changes in fluorescence associated with increases in intracellular Ca^2+^ upon ATP stimulation were measured using confocal microscopy. B.) Representative individual traces of changes in GCaMP6f fluorescence (ΔF/F_0_) in response to ATP stimulation (upper panel) are shown. C.) BAPTA-AM (20 μM, n=5 wells) and CPA (20 μM, n=5 wells) eliminated the ATP (100 μM, n=5 wells)-induced increase in Ca2+ concentration. MAFP (10 μM, n=5 wells) and Bel (10 μM, n=5 wells) did not inhibit ATP (100 μM)-induced increases in Ca^2+^ in astrocytes. ***P≤0.0001, Tukey’s *post hoc* test. All bar graphs show the mean ± s.e.m.

As a broad-based approach to AA-specific lipidomics, Ca^2+^-independent release of AA and/or its metabolites was performed using a [^3^H]-AA assay **(Figure 2A)**. Cultured rat astrocytes were preincubated overnight with [^3^H]-AA with sufficient time to allow for its incorporation into multiple biosynthetic and metabolic pathways. This process includes incorporation into membrane phospholipids, which are precursors for endocannabinoids, and related lipids, which are precursors for AA release (Chen and Chen, 1998; Strokin et al., 2003). Therefore, this assay was used to determine whether any AA precursors or metabolites derived from the [^3^H]-AA incorporated into the cell were being released; hereafter, we refer to the composite of [^3^H]-labeled AA metabolites as [^3^H]-AA. ATP alone failed to induce a detectable release of [^3^H]-AA (**Figure 2B)**; however, in cultures pretreated for 10 to 12 minutes with CPA or BAPTA-AM, ATP induced a robust increase in the release of [^3^H]-AA, whereas CPA alone had no such effect **(Figure 2B)**. Similarly, upon stimulation with a combination of the nonselective mGluR agonist tACPD (100 µM) and the ionotropic glutamate receptor agonist AMPA (100 µM), we observed significant release of [^3^H]-AA when cytosolic Ca^2+^ was blocked with CPA but not in cultures without CPA pretreatment **(Figure 2C)**. These observations confirm that rat astrocytes can release AA derivatives, which are key precursors of bioactive eicosanoids (Strokin et al., 2003; Rosenegger et al., 2015), but that AA release is, surprisingly, inhibited by increases in cytosolic Ca^2+^.

**Figure 2:**
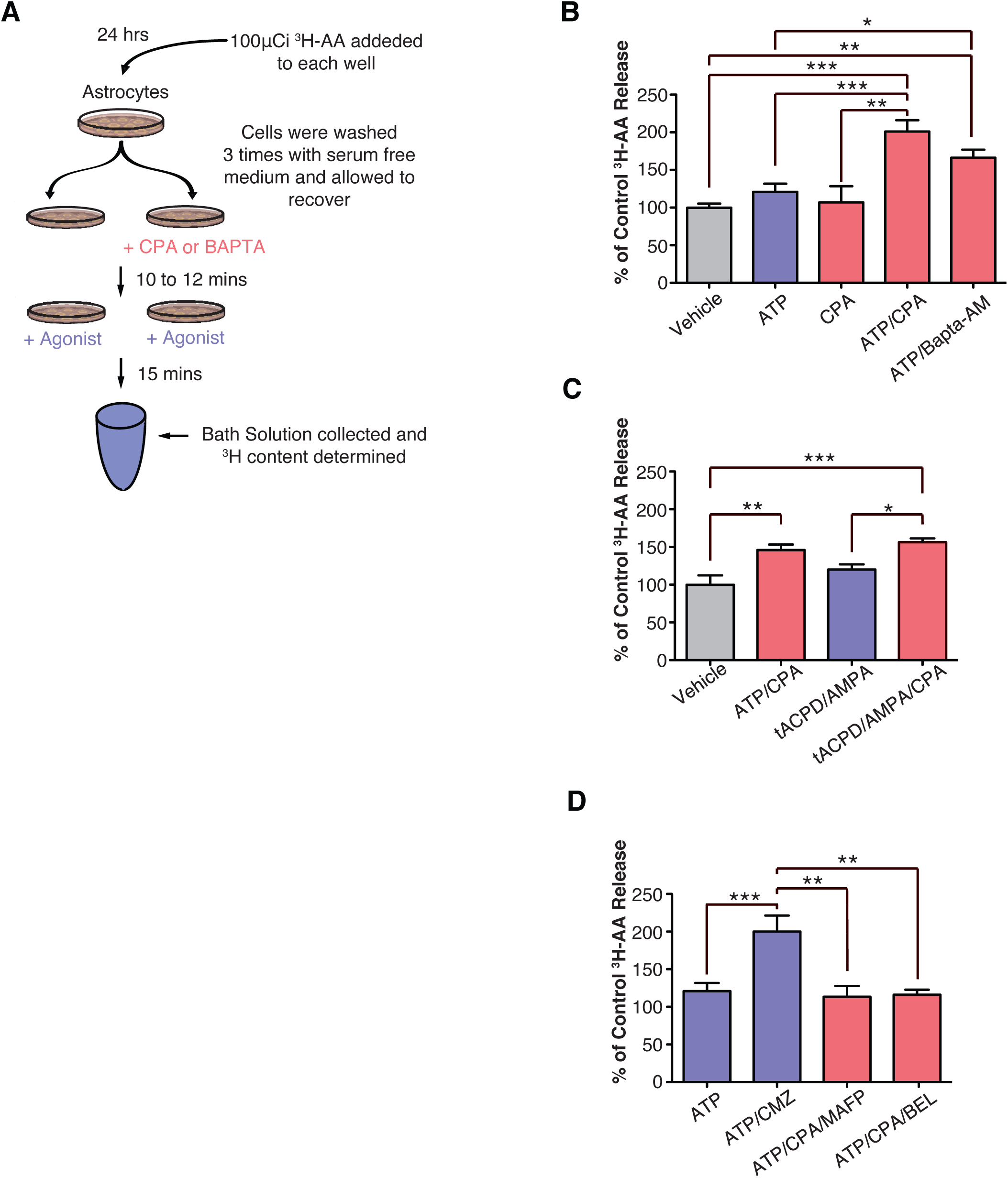
GPCR-mediated Ca^2+-^independent release of ^3^H-AA and/or its metabolites from astrocytic cultures. A.) The schematic shows the ^3^H-AA Radioactive Assay. B.) The effects of ATP (100 μM, n=45 wells), CPA (20 μM, n=9 wells), ATP/CPA (n=44 wells), and ATP/BAPTA-AM (n=24 wells) on Ca^2+^-independent release of [^3^H]-AA and its derivatives in astrocytic cultures were compared to those of the control and each other. ***P≤0.0001; **P≤0.001; *P≤0.05, Tukey’s *post hoc* test. C.) The effects of the mGluR agonists tACPD/AMPA (100 μM, n=7), tACPD/AMPA/CPA (n=7 wells), and ATP/CPA (n=7 wells) on the Ca^2+^-independent release of [^3^H]-AA and its derivatives in astrocytic cultures were compared to those of the control and individual effects. ***P≤0.0001; **P≤0.001; *P≤0.05, Tukey’s *post hoc* test. D.) The effects of the iPLA_2_ inhibitor BEL (10 μM, n=12 wells), the cPLA_2_ inhibitor MAFP (10 μM, n=12 wells), or the calmodulin/Ca^2+^ complex inhibitor CMZ (2 μM, n=20 wells) with ATP on Ca^2+^-independent release of [^3^H]-AA and its derivatives in astrocytic cultures were compared to ATP alone. ***P≤0.0001; **P≤0.001, Tukey’s *post hoc* test. All bar graphs show the mean ± s.e.m.

To explore the mechanism of Ca^2+^-independent release of [^3^H]-AA, we next evaluated whether inhibition of Ca^2+^-sensitive cPLA_2_ or Ca^2+^-insensitive iPLA_2_ enzymes would reduce [^3^H]-AA release. Pretreatment of astrocytes with 10 µM MAFP significantly decreased ATP-induced release of [^3^H]-AA in cultures exposed to CPA **(Figure 2D)**. Similarly, 10 µM BEL treatment suppressed the release of [^3^H]-AA, thus confirming that iPLA_2_ plays a role in Ca^2+^-independent lipid release **(Figure 2D)**. [^3^H]-AA release was observed only when agonist-induced increases in cytosolic Ca^2+^ were blocked **(Figure 2B-C)**, suggesting that intracellular Ca^2+^ inhibits Ca^2+^-independent iPLA_2_.

Calmodulin is a potent Ca^2+^-dependent inhibitor of iPLA_2_ (Wolf and Gross, 1996). To assess the interaction between calmodulin and iPLA_2_, cells were treated with calmidazolium (CMZ), which is an inhibitor of Ca^2+^/calmodulin interactions that has been shown to eliminate calmodulin blocking of iPLA_2_ (Wolf and Gross, 1996). In the presence of CMZ (2 μM), ATP treatment led to the release of a significant amount of [^3^H]-AA, which was comparable to the release induced by blocking increases in cytosolic Ca^2+^ by preloading astrocytes with either CPA or BAPTA **(Figure 2D)**. This observation suggested that Ca^2+^ acts primarily as a brake through calmodulin, which effectively inhibits iPLA_2_ activity (Wolf and Gross, 1996; Wolf et al., 1997). These findings are consistent with previous studies showing that iPLA_2_ is involved in receptor-mediated AA release from pancreatic islet cells (Gross et al., 1993), smooth muscle cells (Lehman et al., 1993), and endothelial cells (Seegers et al., 2002). Taken together, these data provide evidence of a new signaling mechanism in astrocytes involving Ca^2+^-independent iPLA_2_, which is the major PLA_2_ isoform in the brain and accounts for 70% of total PLA_2_ activity (Yang et al., 1999)

### Targeted lipidomics reveals Ca^2+^-independent lipid production in cultured astrocytes

Using lipid extraction and partial purification methods coupled to HPLC/MS/MS, we performed targeted lipidomic screening of cultured astrocytes that were preincubated with CPA and then challenged with ATP and compared the results to that of vehicle control astrocytes. Of the 50 lipids screened, 30 were present in each of the samples and were used for comparative analyses. The lower panel in Figure 3A lists all of the concentrations as the mean ± SEM **(Figure 3A)**. **Figure 3A** summarizes the lipids that showed significant concentration differences, as well as the magnitudes of the differences. Among the 30 lipids detected, 16 (including AA and PGE_2_) increased upon ATP challenge in the presence of CPA. **Figure 3B-C** shows representative chromatograms of the HPLC/MS/MS methods used to detect PGE_2_. Notably, the levels of 5 of the 8 AA derivatives in the set, including docosahexaenoyl ethanolamine, were significantly increased following ATP challenge with CPA preincubation, although the most dramatic increases were observed for the prostaglandin PGE_2_. **Figure 3D** shows the representative differences in concentration of PGE2, DEA, and AA.

**Figure 3.**
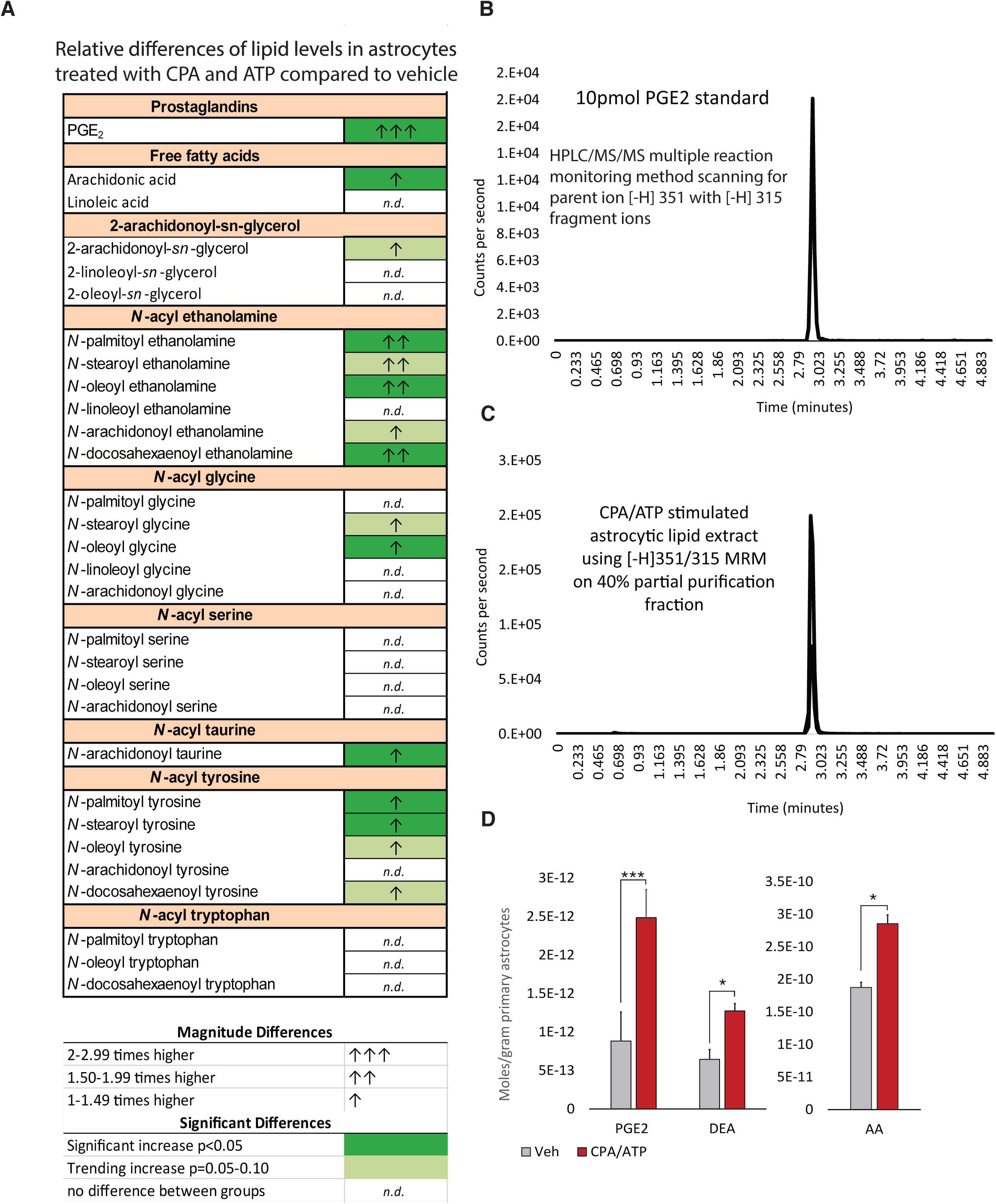
Targeted lipidomics of rat astrocytic cultures using HPLC/MS/MS. A.) The significant differences and the magnitude of change between the vehicle controls and the CPA/ATP-stimulated cells for the 30 lipids that were detected across samples are summarized. Dark green denotes significant differences (p ≤ 0.05); light green denotes increasing trends (0.10 ≥ p ≤ 0.05). No significant difference is indicated by “*n.d*.”. Arrows indicate the magnitude of effects, as indicated in the legend. *The raw data and statistical analyses for all lipids are shown in the figure table.* B.) An example chromatograph of a PGE2 standard (1 pmol) analyzed by the HPLC/MS/MS method that was optimized for this standard in negative ion mode is shown. C.) An example of an analysis of a partially purified astrocytic extract using the HPLC/MS/MS method that was optimized for PGE2 is shown. D.) Differences in the astrocytic production of AA and two of its derivatives, PGE2 and docosahexaenoyl ethanolamine (DEA), were expressed as the average moles/sample; vehicle, n=3 flasks; CPA/ATP, n=4 flasks.

### Ca^2+^-independent release of PGE_2_ from cultured astrocytes

Given that Ca^2+^-dependent lipid release from astrocytes has been previously implicated in vasoregulation (Zonta et al., 2003; Takano et al., 2006; Gordon et al., 2008), we were surprised that PGE_2_ was released via a Ca^2+^-independent mechanism. Our HPCL/MS/MS method requires large quantities of cells to detect PGE_2_, whereas the PGE_2_ ELISA can be used to accurately detect PGE_2_ release in 24-well cultures. Therefore, we used ELISAs to explore the Ca^2+^ dependence of astrocytic PGE_2_ release. Because iPLA_2_ is essential for Ca^2+^-independent liberation of AA-derived lipids, we first assessed whether knockdown of iPLA_2_ via viral transduction would inhibit astrocytic release of PGE_2_. In the presence of CPA (20 μM), ATP (100 μM) failed to induce PGE_2_ release in shRNA-transduced astrocytic cultures, whereas there was a significant increase in the release of PGE_2_ by ATP in the presence of CPA in control shRNA cultures **(Figure 4A)**. Knockdown of iPLA_2_ via shRNA-mediated viral transduction was confirmed by Western blot analysis **(Figure 4B)**.

**Figure 4:**
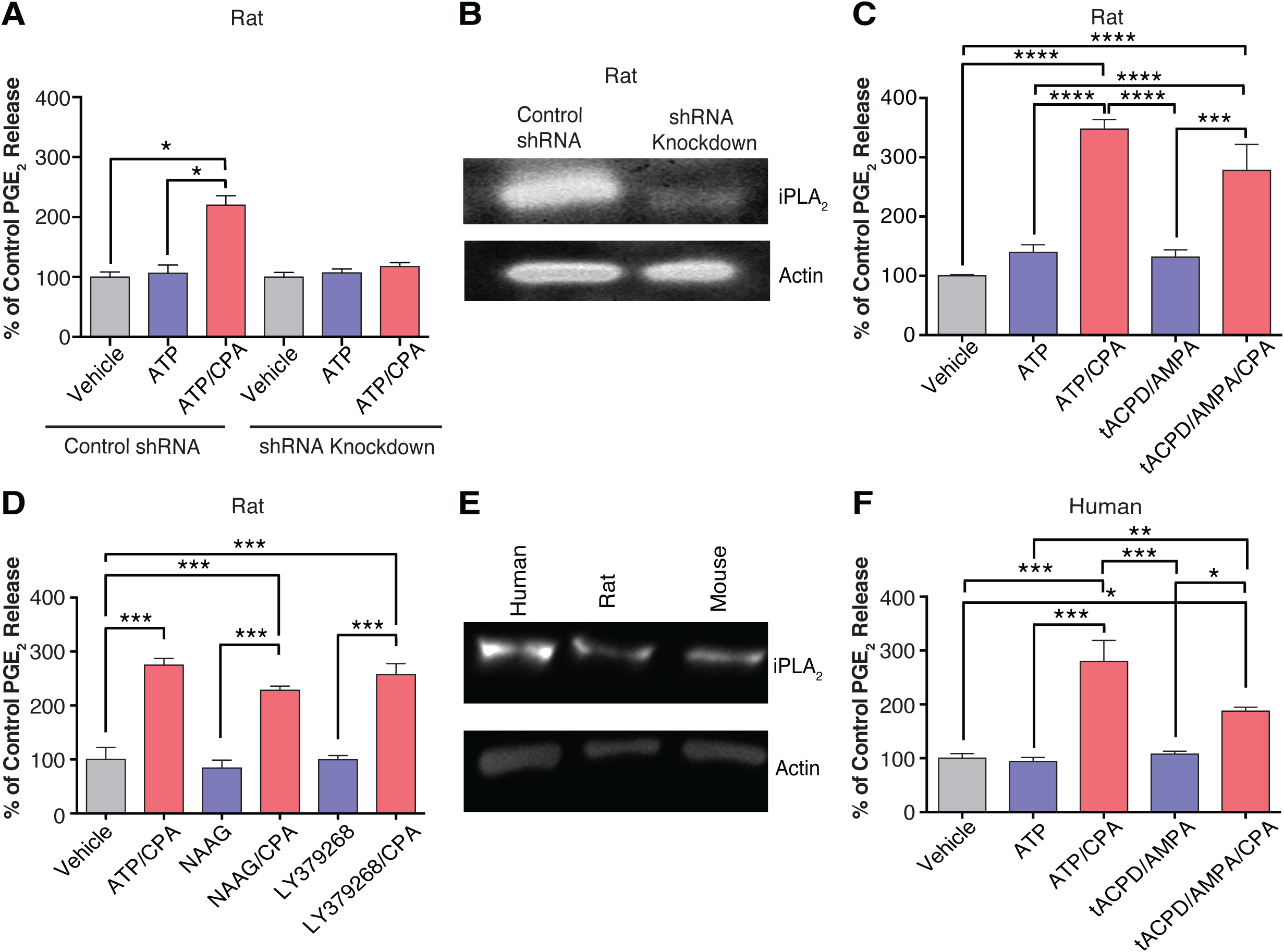
GPCR-mediated Ca^2+^-independent release of PGE_2_ from astrocytic cultures. A.) The effects of ATP (100 μM, n=4 wells) and ATP/CPA (n=4 wells) on Ca^2+^-independent PGE_2_ release in rat astrocytic cultures virally transduced with shRNA to knockdown iPLA_2_ was compared to that in control shRNA cultures. *P≤0.05, Dunn posttest. B.) Western blot analysis of iPLA_2_ knockdown via shRNA was performed. β-actin-specific antibodies were used for the normalization of protein loading. C.) The effects of the GPCR agonists ATP (100 μM, n=12 wells), tACPD/AMPA (100 μM, n=7 wells), tACPD/AMPA/CPA (n=7), and ATP/CPA (n=12 wells) on Ca^2+^-independent PGE_2_ release in rat astrocytic cultures was compared to that in control cultures. ****P≤0.0001; ***P≤0.001, Tukey’s post hoc test. D. The effects of ATP/CPA (100 µM, n=4) and the mGluR3 agonists NAAG (100 μM, n=4 wells), LY379268 (100 μM, n=4 wells), NAAG/CPA (n=4), and LY379268/CPA (n=4 wells) on Ca^2+^-independent PGE_2_ release in rat astrocytic cultures were compared to that in control cultures. ***P≤0.001, Tukey’s post hoc test. E.) Western blot analysis of the iPLA_2_ protein in human, rat, and mouse cultures was performed. β-actin-specific antibodies were used for the normalization of protein loading. F. The effects of the GPCR agonists ATP (100 μM, n=8 wells), ATP/CPA (n=8 wells), tACPD/AMPA (100 μM, n=8 wells), and tACPD/AMPA/CPA (n=8 wells) on Ca^2+^-independent PGE_2_ release in human astrocytic cultures were compared to that in control cultures. ***P≤0.0001; **P≤0.001; *P≤0.05, Tukey’s *post hoc* test. All bar graphs show the mean ± s.e.m.

As cultured astrocytes express mGluR5 (Balazs et al., 1997; Silva et al., 1999; Gebremedhin et al., 2003), we next determined whether the agonists tACPD (100 μM) and AMPA (100 μM) evoked PGE_2_ release in the presence/absence of CPA. In the absence of CPA, these agents evoked little to no release of PGE_2_ **(Figure 4C)**. However, when CPA was used to block the release of Ca^2+^ from internal stores, tACPD and AMPA significantly increased PGE_2_ release (Figure 4C).

The most abundant mGluRRs expressed by astrocytes are mGluR5 and mGluR3 (Petralia et al., 1996; Aronica et al., 2000; Tamaru et al., 2001). However, mGluR5 is developmentally regulated and is not expressed by astrocytes in the adult brain, whereas mGluR3 is persistently expressed at high levels throughout adulthood (Sun et al., 2013). Determining whether the activation of mGluR3 can induce Ca^2+^-independent PGE_2_ release is important because the activation of this receptor has recently been shown to induce Ca^2+^ transients in adult hippocampal astrocytes (Haustein et al., 2014; Tang et al., 2015). In the presence of CPA, the mGluR3 agonists NAAG and LY379268 (Wroblewska et al., 1997; Bond et al., 1999) significantly increased the release of PGE_2_, whereas the same agonists failed to induce the release of PGE_2_ in the absence of CPA **(Figure 4D)**. Notably, this result shows that activation of an astrocytic Gi-coupled receptor can induce the release of gliotransmitters via a Ca^2+^-independent mechanism.

To further investigate human astrocytes, we performed the same experiments in primary cultured astrocytes harvested from human embryonic tissue (Windrem et al., 2004; Windrem et al., 2008; Han et al., 2013). After pharmacologically assessing iPLA_2_ activity in rat astrocytes **(Figure 2D)**, we evaluated iPLA_2_ expression in all culture models. Western blot analysis revealed that iPLA_2_ expression was not limited to rat astrocytes but iPLA2 was also expressed in human and mouse astrocytes **(Figure 4E)**. In the presence of CPA, ATP evoked significant PGE_2_ release from cultured human astrocytes, whereas little to no PGE2 release was observed with ATP alone **(Figure 4F)**. Similarly, coapplication of tACPD and AMPA had the same effect **(Figure 4F)**. Taken together, these findings show that ATP or mGluR3 activation leads to Ca^2+^-independent PGE_2_ release and that iPLA_2_ is expressed in mouse, rat, and human astrocyte cultures.

### Ca^2+^-independent astrocytic lipid release enhances mEPSPs via Kv channel blockade

Thus far, our experiments demonstrated that in response to agonist exposure, cultured astrocytes can indeed release lipids via a Ca2+-independent pathway.- In fact, the in vitro analysis showed that agonist-induced Ca^2+^ increases impeded PGE_2_ release, whereas CPA, BAPTA, and CMZ pretreatment potentiated PGE_2_ release.

Earlier studies have shown that AA and its metabolite PGE_2_ inhibit neuronal Kv currents (Horimoto et al., 1997; Nicol et al., 1997; Evans et al., 1999) and thereby enhance excitability (Sekiyama et al., 1995; Chen and Bazan, 2005; Sang et al., 2005). We therefore evaluated whether astrocytic lipid release also inhibits neuronal Kv currents and enhances neuronal excitability in acute hippocampal slices. We performed dual patch-clamp recordings of pairs of CA1 pyramidal neurons and astrocytes in acute hippocampal brain slices prepared from 12–18-day-old mice **(Figure 5A)**. To isolate transiently active potassium currents, we used 100 ms voltage ramps, which are sufficient to capture transient A-type currents (Phillips et al., 2018). Notably, the I‒V ramp enables one to discern what voltage deflection in the outward current a drug affects (Jackson and Bean, 2007). Kv currents in CA1 neurons are used as an assay for astrocytic lipid release as these currents are sensitive to AA and/or its metabolites (Villarroel and Schwarz, 1996; Carta et al., 2014). We isolated the Kv currents by adding 1 mM QX314 to the patch pipette to block sodium channels (Talbot and Sayer, 1996; Kim et al., 2010) and by imposing a voltage ramp (from -100 mV to 50 mV) every 5 seconds to continuously monitor changes in the Kv current (Ji et al., 2000; Rangroo Thrane et al., 2013b; Carta et al., 2014) to assess agonist-induced astrocytic lipid release (Figure 4A). As expected, direct puffing of PGE_2_ (50 μM) or AA (50 μM) from a micropipette significantly reduced the Kv current, which is consistent with the results of previous studies (Evans et al., 1999; Carta et al., 2014) **(Figure 5B-C)**.

**Figure 5:**
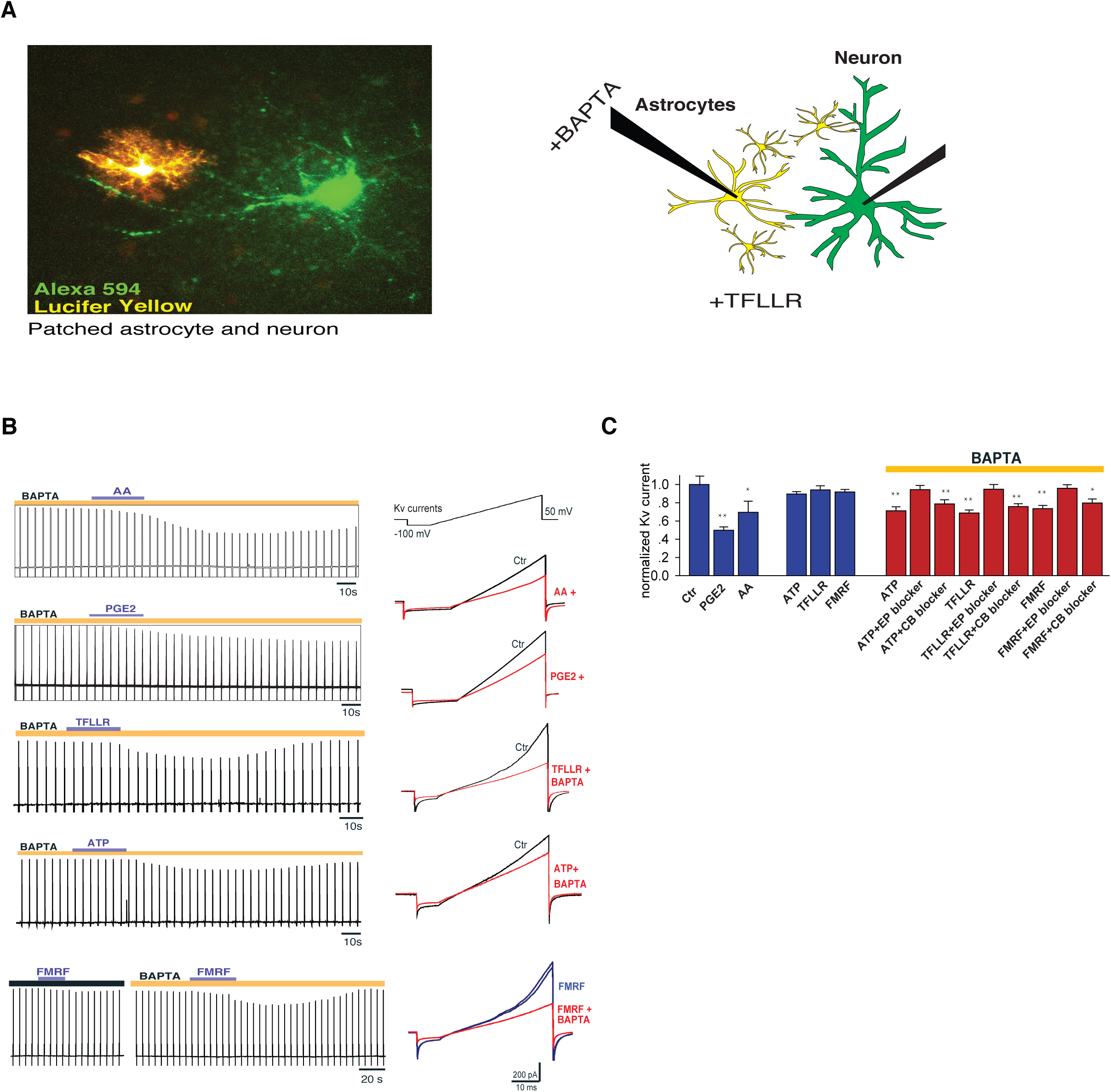
Ca^2+^-independent astrocytic lipid release inhibits Kv blockade. A.) of the image shows a patched astrocyte and a patched neuron filled with fluorescent dyes. B.) Direct application of AA (50 μM) and PGE2 (50 μM) decreased the Kv current. ATP (100 μM) and TFLLR (30 μM) together with BAPTA also decreased the Kv current, as indicated by the peak currents. FMRF (15 μM) decreased the Kv current in MrgA1^+/-^ mice in the presence of BAPTA but not in the absence of BAPTA. Traces to the left are representative recordings of changes in membrane voltages in the presence of the previously described drugs. C.) The effects of different drugs on the voltage ramp-induced voltage-gated K^+^ currents was quantified (**P<0.01; *P<0.05; n =5-7 mice). All bar graphs show the mean ± s.e.m.

To assess the effects of astrocytic Ca^2+^ signaling on neighboring neuronal Kv, we stimulated astrocytes with ATP or TFLLR-NH_2_, an agonist of protease-activated receptor-1 (PAR1), which is primarily expressed by astrocytes (Shigetomi et al., 2008). ATP (100 μM) induced a comparable change in neuronal Kv when BAPTA (50 μM) was present in the astrocyte patch pipette but not in its absence **(Figure 5B-C).** A similar transient decrease in the Kv current was observed when astrocytes were activated by TFLLR-NH_2_ (30 μM). Again, a reduction in neuronal Kv currents was detected only when TFLLR was applied and when BAPTA was added to the astrocytic solution **(Figure 5B-C).** To selectively activate astrocytic Ca^2+^ signaling, we next generated acute hippocampal slices from MrgA1 transgenic animals (MrgA1^+/-^), which express the exogenous Gq-coupled MRG receptor (MrgA1) under the control of the GFAP promoter. The MrgA1 agonist Phe-Met-Arg-Phe amide (FMRF) mobilizes intracellular astrocytic Ca^2+^ stores, thus allowing use to assess the effects of astrocytic Ca^2+^ signaling on neighboring neuronal Kv currents. Although FMRF (15 μM) induces potent and selective increases in astrocytic cytosolic Ca^2+^ in hippocampal slices (Fiacco et al., 2007; Agulhon et al., 2010; Wang et al., 2012), we observed no detectable changes in neuronal Kv current when the pipette used to patch astrocytes did not contain BAPTA **(Figure 5B-C)**. However, when BAPTA was present in the astrocyte patch pipette, there was a marked decrease in the neuronal Kv current evoked by FMRF exposure **(Figure 5B-C)**. Thus, astrocytes can modulate neuronal Kv currents via a previously undocumented Ca^2+^-independent lipid release mechanism.

To assess whether a decrease in the neuronal Kv current is a consequence of Ca^2+^-independent astrocytic lipid release, we employed specific lipid receptor antagonists for PGE_2_ and endocannabinoids. As PGE_2_ receptors 1, 2, 3, and 4 are expressed on hippocampal pyramidal neurons (Andreasson, 2010; Maingret et al., 2017), we used AH6809, a PGE_2_ EP1, 2, and 3 antagonist (Abramovitz et al., 2000; Ganesh, 2014), and GW627368, a PGE_2_ EP4 antagonist (Jones and Chan, 2005; Wilson et al., 2006). With AH6809 (10 μM) and GW627368X (3 μM) in perfusion solution, the TFLLR-, FMRF-, and ATP-induced Ca^2+^-independent decrease in the neuronal Kv current caused by PGE2 was abolished **(Figure 5C)**. In contrast, the CB1 antagonist AM251 (5 μM) failed to abolish the decrease in Kv current **(Figure 5C)**. Taken together, these data suggest that the observed decrease in the neuronal Kv current is a result of astrocytic Ca^2+^-independent release of PGE_2_.

Interestingly, the presence or absence of BAPTA in the astrocytic pipette solution affected agonist-induced changes in neuronal membrane potential **(Figure 6A)**. Without BAPTA in astrocyte pipettes, TFLLR induced hyperpolarization (-2.4 ± 0.26 mV) **(Figure 6A)**, which has been linked to a decrease in extracellular potassium (Wang et al., 2012). However, with BAPTA present in astrocyte pipettes, TFLLR induced depolarization (2.1 ± 0.25 mV) **(Figure 6A)**, an effect that could be attributed to blockage of the potassium current.

**Figure 6:**
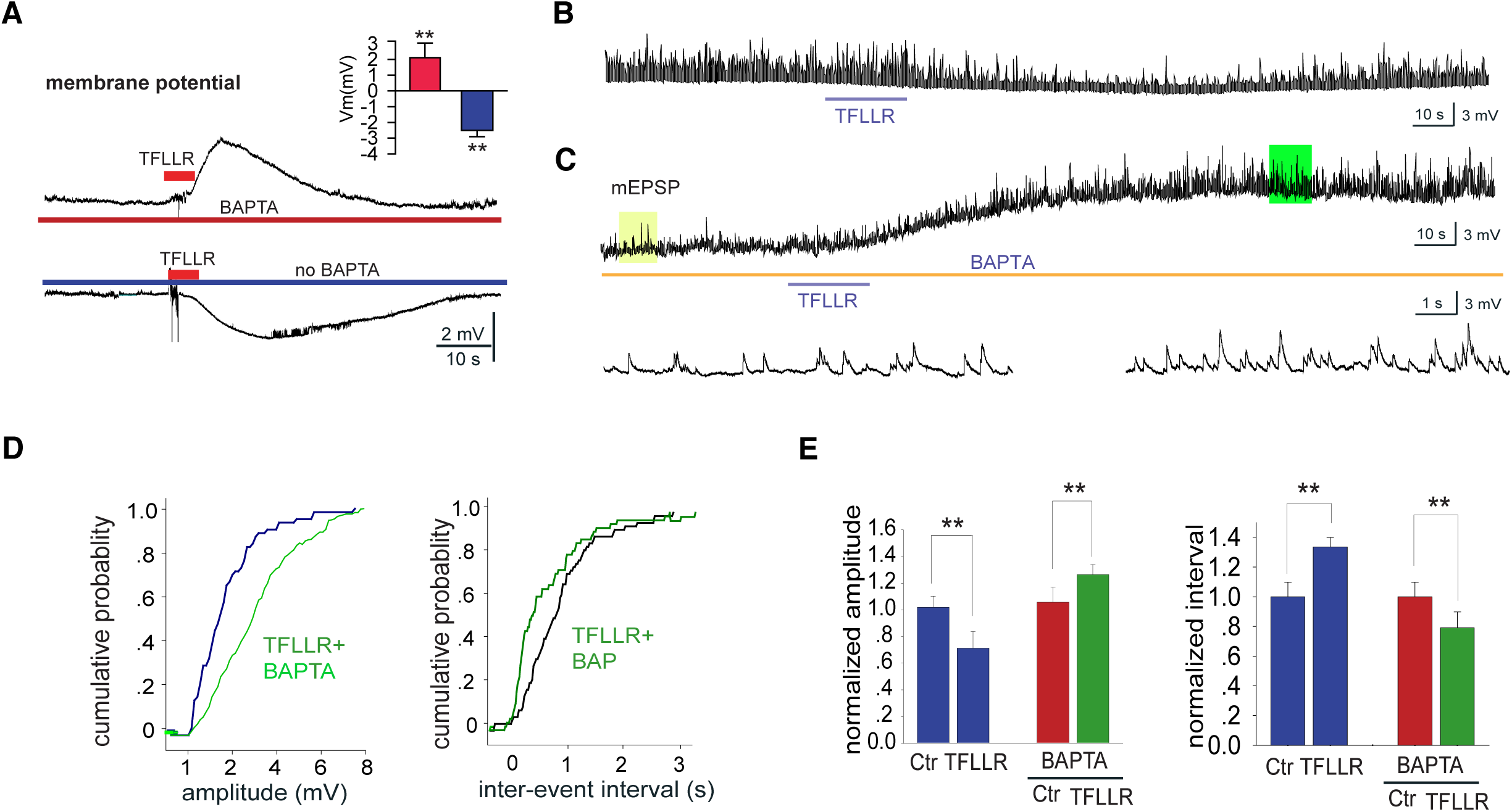
Ca^2+^-independent astrocytic lipid release increases the amplitude and frequency of mEPSP. A.) The representative traces show changes in membrane potential (VM) upon TFLLR application in the presence or absence of BAPTA, and quantitative values are shown in the inset (**P<0.01; t test; n =5-6 mice). B.) The representative trace show hyperpolarization and the reduction in mEPSP amplitude and frequency in excitatory hippocampal neurons upon TFLLR (30 μM) application without BAPTA in neighboring astrocytes. C.) The representative trace shows depolarization and an increase in mEPSP amplitude and frequency in excitatory hippocampal neurons upon TFLLR (30 μM) application with BAPTA present in neighboring astrocytes. The lower traces are extensions of parts of the recordings in the upper traces. D.) The cumulative mEPSP amplitude and cumulative mEPSP frequency distribution recorded in B) were compared. E.) Statistical analysis of the amplitude and interevent intervals before and after TFLLR treatment in the presence and absence of BAPTA was performed (**P<0.001; paired t test; n =5-7 mice). All bar graphs show the mean ± s.e.m.

PGE_2_ has been shown to enhance neuronal mEPSPs (Sekiyama et al., 1995) (Sang et al., 2005). We found that TFLLR-induced activation of astrocytes triggered a decrease in mEPSP amplitude and frequency in acute hippocampal slices from normal mice **(Figure 6B)**. The opposite effect, an increase in the amplitude and frequency of mEPSCs, was observable only when BAPTA was added to the astrocytic pipette solution **(Figure 6C)**. This observation supports the notion that astrocytic Ca^2+^-independent lipid release may function as a signaling mechanism that can modulate proximal synaptic activity by blocking Kv, which in turn may increase the frequency and amplitude of mEPSCs. Interestingly, the opposing effects of TFLLR on mEPSP amplitude and frequency were dependent upon astrocytic Ca^2+^ levels (**Figure 6D-E**), suggesting that Ca^2+^ depletion provides a pathway that enhances neuronal excitability by astrocytic PGE2.

## DISCUSSION

Lipidomics has the potential to open exciting new avenues within the field of gliotransmission. In the present study, we utilized a series of lipidomic methodologies to show that Ca^2+^ chelation followed by metabotropic glutamate or purinergic receptor stimulation promoted the formation of a variety of lipids (**Figures 2, 3**) in astrocytes and drove the release of PGE_2_ in rat and human embryonic astrocyte cultures (**Figure 4**). In addition, we demonstrated that receptor-mediated Ca^2+^-independent PGE_2_ release modulates neuronal Kv, resulting in enhanced synaptic activity in slices, as was determined by the increase in mEPSP amplitude and frequency **(Figures 5, 6)**. Multiple findings presented herein also support our hypothesis that this agonist-induced and Ca^2+^-independent lipid release from astrocytes depends on the activation of iPLA_2_. These observations represent, to our knowledge, the first evidence of Ca^2+^-independent release of lipid gliotransmitters and thus add a novel dimension to our understanding of glial–neuronal communication.

### Importance of the iPLA_2_ pathway in Ca^2+^-independent astrocytic lipid production

Although astrocytes are known to constitutively release AA and other fatty acids (Moore, 2001; Bouyakdan et al., 2015), the pathways involved in constitutive AA release are currently not known; however, inverse agonists of purinergic receptors (Ding et al., 2006) and mGluR (Carroll et al., 2001) might serve as useful pharmacological tools for investigating constitutive PLA_2_ activity in relation to gliotransmission. To date, many studies have shown the multifaceted functions of PLA_2_ in the CNS, but iPLA_2_ accounts for 70% of the PLA_2_ activity in the rat brain (Yang et al., 1999). Although the effects of cPLA_2_ activation are well documented (Malaplate-Armand et al., 2006; Schaeffer and Gattaz, 2007; Kim et al., 2008), iPLA_2_ has also been shown to participate in phospholipid remodeling (Sun et al., 2004) and regulate hippocampal AMPA receptors involved in learning and memory (Menard et al., 2005). Furthermore, iPLA_2_ regulates store-operated Ca^2+^ entry in cerebellar astrocytes (Singaravelu et al., 2006) and provided neuroprotection in an oxygen–glucose deprivation model (Strokin et al., 2006). Astrocytes express both isoforms of PLA_2_ (Sun et al., 2005), and numerous studies have reported that the release of AA and its metabolites is under the regulation of Ca^2+^-dependent PLA_2_ (cPLA_2_) signaling in astrocytes (Bruner and Murphy, 1990; Stella et al., 1994; Stella et al., 1997; Chen and Chen, 1998). Previous studies have demonstrated astrocytic release of DHA via iPLA2 (Strokin et al., 2003, 2007). Here, we focused on determining whether the largely unexplored iPLA_2_ lipid pathway mediates PGE2 release from astrocytes. Given the ubiquitous expression of iPLA_2_ in astrocytes throughout the brain, we contend that various receptor-activated pathways rely on iPLA_2_ for the regulation of lipid release from astrocytes **(Figure 7)**.

**Figure 7:**
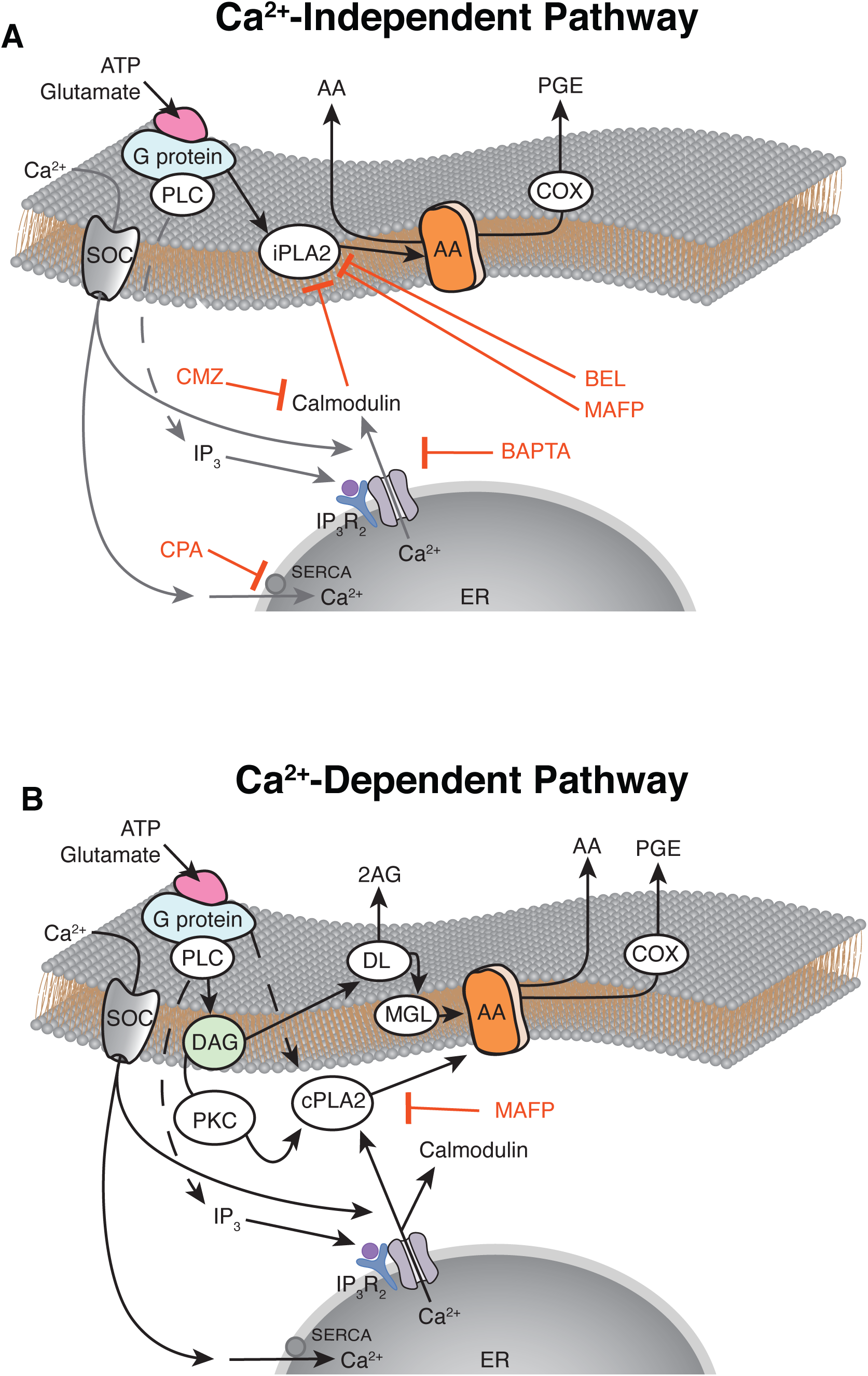
Models of GPCR-mediated Ca^2+^-independent and Ca^2+^-dependent lipid release in astrocytes. A.) The schematic depicts receptor-mediated Ca^2+^-independent lipid release across the astrocytic membrane. B.) The schematic depicts receptor-mediated Ca^2+^-dependent lipid release across the astrocytic membrane.

### Ca^2+^-independent astrocytic PGE_2_ release rapidly modulates neuronal Kv

An important aspect of astrocytic gliotransmitter release is timing. Agonist-induced increases in astrocytic Ca^2+^ occur on a slow time scale of seconds (Cornell-Bell et al., 1990; Wang et al., 2006; Srinivasan et al., 2015), which allows for the integration of many factors on that time scale. Increased intracellular Ca^2+^ levels are key in the release of gliotransmitters (Parpura et al., 1994; Bezzi et al., 1998; Kang et al., 1998), such as ATP (Coco et al., 2003; Parpura and Zorec, 2010; Illes et al., 2019) and D-serine (Mothet et al., 2000; Yang et al., 2003; Li et al., 2018; Neame et al., 2019). We have recently shown that agonist-induced astrocytic Ca^2+^ signaling can modulate synaptic activity by promoting K^+^ uptake, resulting in a transient decrease in extracellular K^+^ and a decrease in synaptic activity (Wang et al., 2012). More recently, we demonstrated that astrocytes can modulate rapid synaptic activity (≥ 500 ms) via 2AG release upon agonism of mGluR3 receptors (Smith et al., 2019). Because receptor-mediated astrocytic Ca^2+^ drives K^+^ uptake and the release of gliotransmitters, these processes must occur over a relatively prolonged time course (>500 ms). However, agonist-induced Ca^2+^-independent iPLA_2_ lipid release does not require the mobilization of intracellular Ca^2+^ stores; thus, a much faster time scale for signaling, within 10 seconds to milliseconds, is possible. We speculate that iPLA_2_-mediated lipid release may act as a feedback system that enhances fast synaptic transmission in the short term or under conditions of minimal neuronal activity, whereas subsequent activity-mediated increases in Ca^2+^ serve as a brake in the form of calmodulin-dependent inhibition of iPLA_2_. The slower Ca^2+^-dependent release of gliotransmitters and stimulation of K^+^ uptake seems more suited toward the slow and widespread modulation of brain activity (typically inhibition) that occurs in the setting of, for example, tonic activation of the locus coeruleus and consequent norepinephrine release (Bekar et al., 2008; Ding et al., 2013).

Voltage-gated potassium channels are located on the dendrites of hippocampal pyramidal neurons (Johnston et al., 2000), where they play a major role in controlling dendritic excitability by modulating the amplitude of EPSPs. A morphological study revealed that the density of Kv in the dendrites of pyramidal neurons increased 5-fold from the soma to the most distal point in apical dendrites (Hoffman et al., 1997). Inhibition of voltage-gated K^+^ currents consequently enhances EPSPs, possibly explaining why PGE_2_ enhances synaptic transmission and increases LTP in the hippocampus (Sang et al., 2005).

### Lipidomics models as a guide for future gliotransmitter discoveries

Lipidomics is one of the fastest-growing branches in the field of metabolomics and offers the possibility of describing the enormous diversity of lipid species throughout the body, especially in the brain, which is largely composed of lipids (Sinclair, 1975). The complex and fine structure of astrocytic processes gives astrocytes a larger surface area-to-volume ratio than most other types of cells, which makes them uniquely responsive to changes in the extracellular milieu. HPLC/MS/MS lipidomics techniques have provided a means to quantify specific lipid species with an accuracy and sensitivity that is not possible with more traditional methods employing radiolabeled fatty acids and ELISA. Here, we combined these classical techniques with lipidomics to test the hypothesis that lipids are produced and released from astrocytes in a Ca^2+^-independent manner. Although our present focus was on the release of PGE_2_, we found increases in the release of 15 additional lipids. Notably, among the 30 lipids measured in the targeted HPLC/MS/MS assay, the levels of 14 did not change upon the activation of glial receptors, supporting of the idea that GPCR-stimulated, Ca^2+^-independent lipid release is specific. Interestingly, two of the lipids released in response to agonist exposure, namely, *N*-arachidonyl taurine and *N*-palmitoyl tyrosine, were recently shown to activate TRPV4, a cation channel involved in osmotic sensitivity and mechanosensitivity (Raboune et al., 2014). Emerging evidence suggests that the TRPV4/AQP4 complex regulates the response to hypo-osmotic stress in astrocytes (Benfenati et al., 2011). The lipidomics data presented here thus elucidated unpredicted signaling mechanisms that involve astrocytic-derived lipid modulators **(Figure 3)**.

### Potential physiological relevance of calcium-independent lipid release

Astrocytes are known to respond to single experimental stimulation events with small calcium increases in their distal fine processes (Panatier et al., 2011). This effect results from the synchronous firing of as many as 50-1000 synapses in the astrocyte microdomain, as may occur when a large stimulation is applied as far as 500 µm away. However, the spontaneous activity of asynchronous synaptic events does not need to elicit calcium events in astrocytes. Metabotropic GluR- and AMPA-mediated activation of astrocytes may elicit iPLA_2_ activation and the concerted release of AA metabolites and PGE_2_ to affect pre- and postsynaptic potassium channels on nearby neurons. When the stimulation frequency and/or amplitude increase, increases in astrocytic calcium then serve to suppress the overactivation of synapses. In this way, astrocytes may help maintain the strength of relatively quiescent synapses in a calcium-independent fashion while employing multiple calcium-dependent mechanisms to suppress overactivation. Substantiation of this model will require rigorous testing of the spatial and temporal dynamics of astrocytic and individual dendritic activation but is limited by current methodological capabilities that do not allow for electrical recording of individual dendritic spines.

In conclusion, Ca^2+^-independent astrocytic lipid release constitutes a largely unexplored factor in the regulation of complex neuro-glial signaling interactions. Our present analysis contributes to the understanding of agonist-induced Ca^2+^ signaling by demonstrating that the agonism of several Gq and Gi-linked astrocytic receptors can promote the release of lipid modulators and that increases in cytosolic Ca^2+^ act as a brake to prevent PGE_2_ release.

## Author contributions

F.W., H.B.B., and N.A.S. performed the experiments. F.W., H.B.B., J.X.,S.C., and N.A.S. analyzed the data. S.P. and B.J. performed the Western blotting. S.G. and B.L. made the cultures. L.B. and N.A.S. planned the experiments. D.C.M. provided the human astrocytes. F.W., H.B.B., and N.A.S. wrote the paper.

## Acknowledgments

We thank Dr. Ken McCarthy for generously sharing the transgenic mice. This work was supported by the National Institutes of Health Grants K01NS110981 and NSFNCS-FR 1926781 and the Department of Defense Army Research Office Award W91NF2020189 to N.A.S. We thank Vittorio Gallo, Baljit Khakh, Bartosz Kula, Stefano Vicini, Alexander S. Thrane, Vinita Rangroo Thrane, Takahiro Takano, Paul Cumming, and Fernando R. Fernandez for comments and critical discussion on this manuscript. The authors declare no competing financial interests.

